# Unveiling early visual cortical mechanisms in perceptual filling-in: a parametric study of eccentricity and movement dependence

**DOI:** 10.1101/2025.11.11.687787

**Authors:** Kenshu Koiso, Anna Razafindrahaba, Vincent van de Ven, Federico De Martino, Mark J. Roberts, Peter De Weerd

## Abstract

Surface perception relies on interactions between boundary encoding and surface filling-in. In a perceptual filling-in paradigm, a blank figure becomes perceptually replaced by a textured background after prolonged fixation of a point away from the figure. Filling-in begins when neurons representing the figure’s boundary adapt, allowing background-related activity to spread into the figure’s retinotopic representation. Adaptation proceeds faster the better the boundary is stabilized in a neuron’s receptive field (RF). We hypothesized that moving the figure boundary beyond a neuron’s RF would reduce adaptation and hinder filling-in, with larger movements permitting progressively less filling-in. As RF size increases with eccentricity, we further hypothesized that greater eccentricities would require larger movements to interfere with filling-in. Our results confirmed both predictions. The reduction in filling-in duration with increasing motion range permitted estimating RF size at each eccentricity. The slope of a linear function relating RF size to eccentricity matched values reported in human fMRI studies of V1/V2, suggesting that boundary adaptation involves early visual areas. We also explored whether microsaccade amplitude affects filling-in, but found no supporting evidence. Thus, external figure motion and microsaccades may disrupt adaptation through different mechanisms. These findings provide new insights into neural adaptation processes preceding perceptual filling-in.

## Introduction

During the processing of visual images, information that the retina and LGN send to the visual cortex mainly represents local contrasts corresponding to image boundaries (Corbett and Chen, 2018; Hubel and Wiesel, 1959; Wiesel and Hubel, 1963). Therefore, the perception of homogeneous surfaces may require neural interpolation processes in retinotopically organized visual cortex, initiated at the boundary representations (Grossberg, 2003; Neumann et al., 2001; Paradiso et al., 1989; Roe et al., 2005). Several theoretical proposals suggest that boundaries are crucial in both initiating and containing lateral neural spread of activity that might underlie the perceptual filling-in of surfaces (Anstis, 2010; De Weerd, 2006; Pessoa et al., 1998; Weil and Rees, 2011). Moreover, anatomical and functional compartments within low-level visual areas (Sincich and Horton, 2005) as well as computational modeling (De Weerd, 2006; Grossberg and Todorovic, 1988; Neumann et al., 2001) suggest distinct but interacting systems for surface and boundary representation.

One way of studying the interaction between boundary-related processes and the perceptual filling-in of surfaces is the Troxler fading paradigm (Troxler, 1804). In our implementation of this paradigm, a blank figure in a dynamic texture image perceptually fades after a period of maintained fixation (De Weerd et al., 1995; Hunzelmann and Spillmann, 1984; Welchman and Harris, 2001).

According to the two-stage model of perceptual filling-in (De Weerd et al., 1998), neural adaptation of figure boundary representations precedes lateral neural spread of surface features from the background into the topographic representation of the figure (De Weerd et al., 1995) via horizontal connectivity. The degree to which the figure is stabilized on the retina is a major factor in determining the delay between stimulus presentation and perceptual filling-in. In the case of perfect stabilization, perceptual filling-in occurs within about 100ms (Paradiso and Nakayama, 1991), whereas under natural fixation conditions, which implies less perfect retinal stabilization, several seconds of maintaining gaze is required to induce perceptual filling-in (De Weerd et al., 1998; Ramachandran et al., 1993; Spillmann and Kurtenbach, 1992). In addition, during natural fixation, the onset of perceptual filling is preceded by periods of reduced microsaccades (MS; typically eye movements of < 1°), while filling-in offset is related to the occurrence of MS (Martinez-Conde et al., 2004). These findings suggest that neural responses to the figure boundaries show more adaptation the better the retinal image is stabilized, and, correspondingly, an earlier onset of perceptual filling-in.

Neuronal adaptation occurs when a stimulus is maintained in the receptive field (RF) of a neuron for a prolonged period, whereas sufficient movement of the stimulus, or of the gaze position, shifts the stimulus to new un-adapted RFs. Thus, to prevent adaptation, movement of the stimulus or gaze by at least the width of the RF must be sufficient (Fig. 1A). The relationship between movement and adaptation thus provides insight into the size of the relevant RF size and of the cortical area, as RF size increases along the cortical hierarchy (Dumoulin and Wandell, 2008; Wandell and Winawer, 2015). Support for the idea that early-level cortical neurons with small RFs are the primary contributors to boundary representations, adaptation and filling-in, may already be found in demonstrations that MSs interfere with filling-in (Martinez-Conde et al., 2006; Troncoso et al., 2008), as the small size of MSs has been suggested to imply the importance of small RFs, characteristic of early visual cortex. However, the effectiveness of MSs in preventing filling-in may be due to one of several neural processes related to MSs, only one of which is the movement of the figure over the retina (Herrington et al., 2009). For example, MSs may cause saccadic suppression (Bosman et al., 2009; Greilich et al., 2024; Lowet et al., 2016; Matin, 1974; Zuber and Stark, 1966), or feedback signals entering low-level visual cortex (corollary discharge) (Sperry, 1950). These feedback signals may disrupt filling-in independently of the extent of the shift in retinal image. Thus, it is not fully established that the shift in retinal image caused by MSs is the primary cause of their disruptive effect on perceptual filling-in.

**Figure 1.**
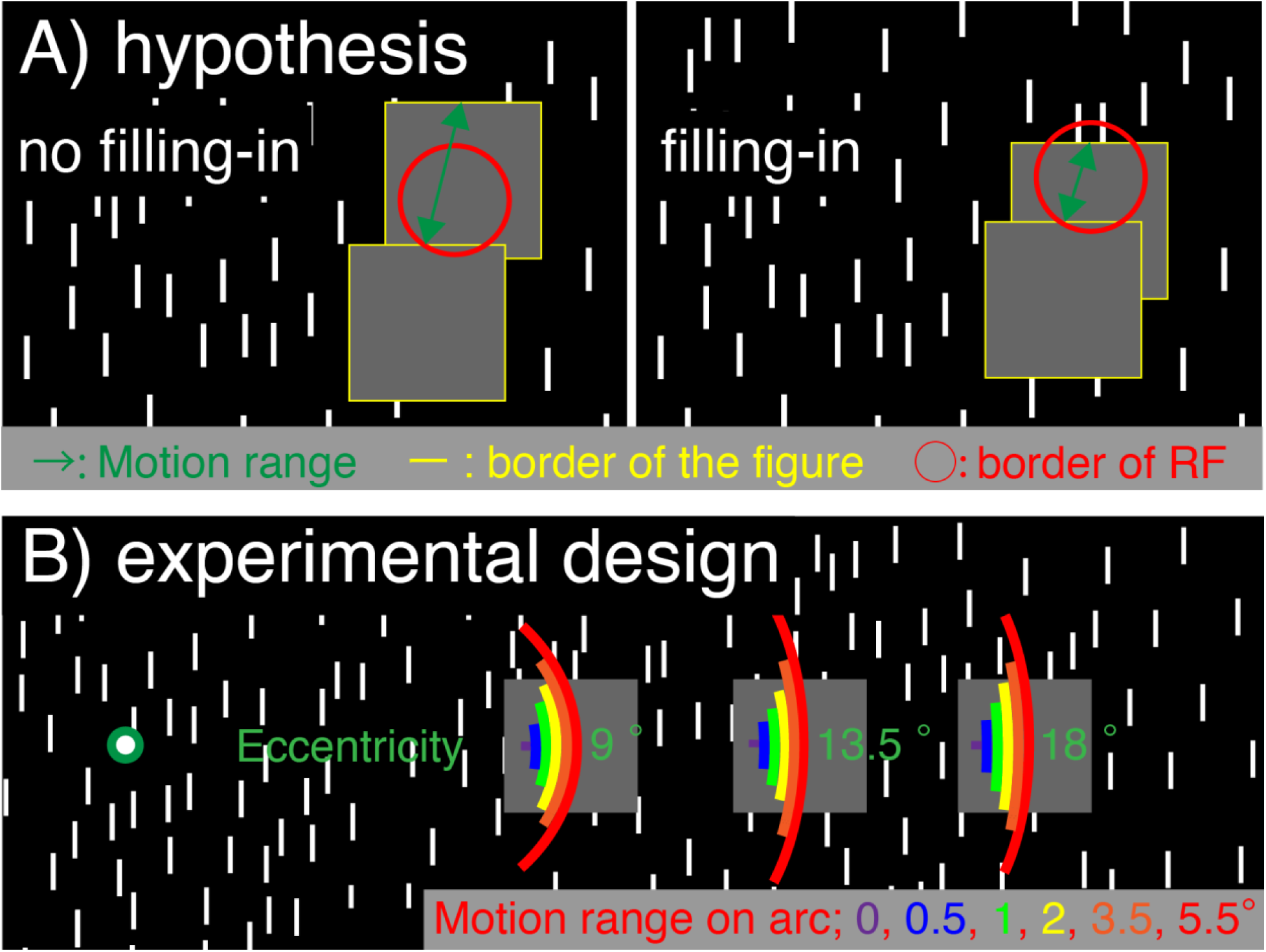
Stimulus and hypothesis. **A)** Filling-in stimulus made of a dynamic texture of vertical line elements on a dark background in which frames were refreshed at a rate of 15 Hz. The figure (grey patch) was isoluminant to the mean luminance of the background. A motion range (green arrow) bringing the edge of the figure (yellow line) from within the RF of a neuron (red outline) to outside the RF (and vice versa for the opposite motion) is hypothesized to interfere with filling-in, as this would counteract adaptation. A motion range small enough to keep the edge of the figure within the RF of a neuron is hypothesized to interfere less with filling-in, as it would still permit sufficient adaptation of the neuron. **B)** The figure was presented at eccentricities of 9°, 13.5°, or 18°. At all three eccentricities, the figure was moved along an equi-eccentric arc, with lengths ranging from 0 to 5.5° (details in Methods). At the 9° eccentricity, an additional set of arc lengths was used, with lengths ranging from 0 to 3.7° (details in Methods).

In this study, we investigated effects of movement of the boundary on filling-in. We experimentally moved the figure back-and-forth over ranges of different distances across the texture background while participants maintained fixation (Fig. 1B) to estimate the size of figure movement that was sufficient to suppress perceptual filling-in. To further assess the critical RF size and cortical areas, we manipulated the eccentricity of the filling-in figure. RF size (expressed as the square root of its surface) increases linearly with increasing eccentricity in early and mid-level visual areas (Dumoulin and Wandell, 2008; Gattass et al., 1987), and the slope with which RF sizes increase as a function of eccentricity differs among visual cortical areas (Dumoulin and Wandell, 2008; Gattass et al., 1988, 1981; Lima et al., 2023; Wandell and Winawer, 2015). Therefore, by measuring the increase with eccentricity in the movement still permitting perceptual filling-in, we reasoned that the slope of the maximum movement for filling-in expressed as a function of eccentricity, would be informative in determining the hierarchical level of areas contributing to boundary adaptation phase of perceptual filling-in. Additionally, we compared the effect of small and large MSs on perceptual filling-in. If the primary mechanism by which MSs disrupt perceptual filling-in is via figure boundaries remaining in the confines of RFs, then at a given eccentricity, larger MSs might be more effective in disrupting adaptation than small MSs.

In summary, the primary aim of the present study was to use a psychophysical approach to assess the neural mechanisms contributing to adaptation prior to perceptual filling-in. First, we aimed to estimate the hierarchical level of visual cortical contributions to and perceptual filling-in. Second, we compared the effects of retinal motion induced experimentally to the effect of retinal motion induced by MSs.

## Methods

### Participants

14 healthy participants (10 females, age range of 18-33 years old) with normal or corrected-to-normal vision participated in this study. One was the first author. Five participants attended all four sessions, four participants attended three sessions, and three participants attended two sessions. Most participants were recruited from the Psychology Bachelor program and its staff at Maastricht University. Participants had no history of neurological disorders. Participants provided informed consent and received study credits or monetary compensation in exchange for participation. Ethics approval was obtained from the Ethics Review Committee for Psychology and Neuroscience (ERCPN, OZL_261_148_12_2022), following the principles expressed in the Declaration of Helsinki.

### Stimuli and Experimental Design

Participants sat with their heads in a chin-and-head rest at a fixed distance of 57 cm from an Iiyama monitor (field of view of size 38 x 30 cm, resolution 1280 x 1024 pixels, refresh rate 60 Hz). The display had a size of 36.87° x 29.49° and was filled with a dynamic texture field except for an isoluminant grey square figure placed either to the left or right from a centrally placed fixation spot (counterbalanced over sessions). The texture was created by randomly setting non-overlapping white bars (0.06° x 0.72° - 2 x 24 pixels) on a black background with a ratio of white bars to black background of 2.66 %. The dynamic texture field was refreshed at 15 Hz by pseudo-randomly selecting frames from 25 random versions of the texture while preventing immediate repetitions of the same pattern. The figure subtended 2.5° x 2.5° and was equi-luminant to the mean luminance of the texture (35 cd/m^2). The fixation spot, a white spot surrounded by a green ring, was presented in the center of the screen.

We tested filling-in with figures placed at three eccentricities from the fixation spot (9°, 13.5°, and 18°, see Fig. 1B), in three separate sessions. At each eccentricity, we tested how moving the figure influenced filling-in with multiple motion ranges (MR). The figure moved in an arc with lengths of 0°, 0.5°, 1°, 2°, 3.5°, or 5.5°, staying equidistant from the fixation spot. Additionally, in a fourth session, we also tested filling-in at 9° eccentricity using a set of shorter motion arcs: 0°, 0.33°, 0.67°, 1.33°, 2.33°, and 3.67°. Each motion range condition was presented three times per block in random order (18 trials per block). Seven blocks were presented per session (126 trials), yielding 21 repetitions per motion condition per session.

The frequency of the motion was constant (4.5 movement cycles per 25 sec trial) across conditions. This was achieved by varying the average speed of each condition (0.18 deg/sec, 0.36 deg/sec, 0.72 deg/sec, 1.26 deg/sec and 1.98 deg/sec for 0.5°, 1°, 2°, 3.5° and 5.5° motion range conditions respectively). To avoid sudden onsets and offsets of motion, the speed of the motion varied along the path according to a sine wave, with higher speed across the center of the path, slowing to 0 at the ends before reversing direction.

### Experimental procedures

#### Eye-data recording

Eye movements and pupil size of the right eye were tracked using the pupil-glint vector at a sampling rate of 225.4 Hz using a ViewPoint eye-tracker (Arrington Research, Scottsdale, AZ) and ViewPoint software version 2.9.5.128. Calibration was performed using the inbuilt 9 points calibration procedure at the beginning of each session. It was complemented by an additional calibration task at the beginning of each block (Nyström et al., 2013), in which five spots (center of the screen and four corners of 3.5° square located at the center of the screen) were presented sequentially. Participants were instructed to direct their gaze to each spot and to maintain fixation while holding the space bar key until the spot disappeared, after which the spot was presented in the next location. The calibration task repeated one time per location per block.

#### Perceptual Filling-in task

Each filling-in trial started with a grey screen containing the fixation spot. Participants pressed and released the spacebar once they had fixated their gaze on the fixation spot, which triggered a 25 sec long stimulus presentation. During this 25 sec stimulus presentation, participants signaled the start of an episode of perceptual filling-in by holding down the space bar on the keyboard in front of them for as long as they experienced perceptual filling-in and releasing the bar when the figure perceptually re-appeared.

### Analysis

#### Behavioral data analysis

To assess filling-in for each motion range condition, we calculated the total filling-in duration in each condition. This involved summing the duration of all filling-in episodes to compute the total filling-in time per trial. We then calculated the means across all trials for each condition for each participant. The analysis of differences among conditions in the total filling-in duration was based on these means per participant. Statistical test was done using repeated measures analysis of variance (ANOVA) or t-test in MATLAB R2023a. In case of non-significant differences for ANOVA or t-test, we calculated Bayes Factors with default prior, Cauchy prior of 0.707 using a publicly available library in matlab (Bart Krekelberg, 2024) or JASP (version 0.95.4) by JASP Team (2025). We then used Jeffrey’s scale to interpret the Bayes Factors (Jeffreys, 1998).

We quantified the limit of the figure motion range of effective filling-in as the motion range that still allowed 50% of duration of filling-in (MR50) relative to the filling-in duration in the no-motion (0°) condition. To estimate the MR50, we normalized the total filling-in duration by the total filling-in duration in the no-motion control condition for the corresponding eccentricity for each eccentricity and each motion condition for each participant. After normalization, we fit an exponential function to the data plotted as a function of motion range and determined the MR50 from the fitted function, using the “fit” function in MATLAB R2023a. The form of the fitted function was:

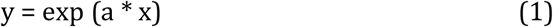

Where y corresponds to the normalised total filling-in, x to the motion range, and a to the exponential factor that was fit to our data.

#### Eye-data processing

The eye-data was processed in MATLAB R2023a. Eye-data recordings were extracted for the full length of the 25 sec filling-in trials and downsampled to 100 Hz to reduce necessary computing power. To detect MSs, we applied the Engbert and Kliegl (2003) MS detection algorithm. We used smoothed horizontal and vertical eye signals over 3 data samples and a velocity threshold criterion of 4 standard deviations above the median velocity (Engbert and Kliegl, 2003) to classify a signal as a MS. Due to technical issues, eye data was not recorded for two participants at the eccentricity of 13.5°. Therefore, MS analysis was limited to only eight participants at that eccentricity. MS analysis in all other eccentricity conditions was performed on all 10 participants.

## Results

We investigated effects of eccentricity and motion range on total filling-in duration. Notably, we found that while moving the figure reduced filling-in, the reduction was gradual, and that substantial filling-in still occurred for moving stimuli, especially for the higher eccentricities (Fig. 2A). To test these effects, we first performed a repeated measures ANOVA with factors eccentricity and motion range, with participant ID as a random factor. We found that filling-in significantly decreased as a function of motion range and increased as a function of eccentricity (eccentricity, F(2, 162) = 16.2, p < 0.001; motion range, F(5, 162) = 44.6, p < 0.001; interaction, F(10, 162) = 3.04, p = 0.0015). To further quantify the relationship between filling-in and motion range, we normalized total filling-in duration for each condition by the total filling-in observed in the 0° motion range condition for each eccentricity separately. The motion range x eccentricity interaction and main effect of motion range were both highly significant (interaction, F(10, 162) = 4.1, p < 0.001; motion range, F(5, 162) = 88.7, p < 0.001). There was no significant main effect of eccentricity (F(2, 162) = 0.07, p =0.928), in line with the normalization per session. This analysis indicated a steeper decrease in filling-in as a function of motion range at the smaller eccentricities.

**Figure 2.**
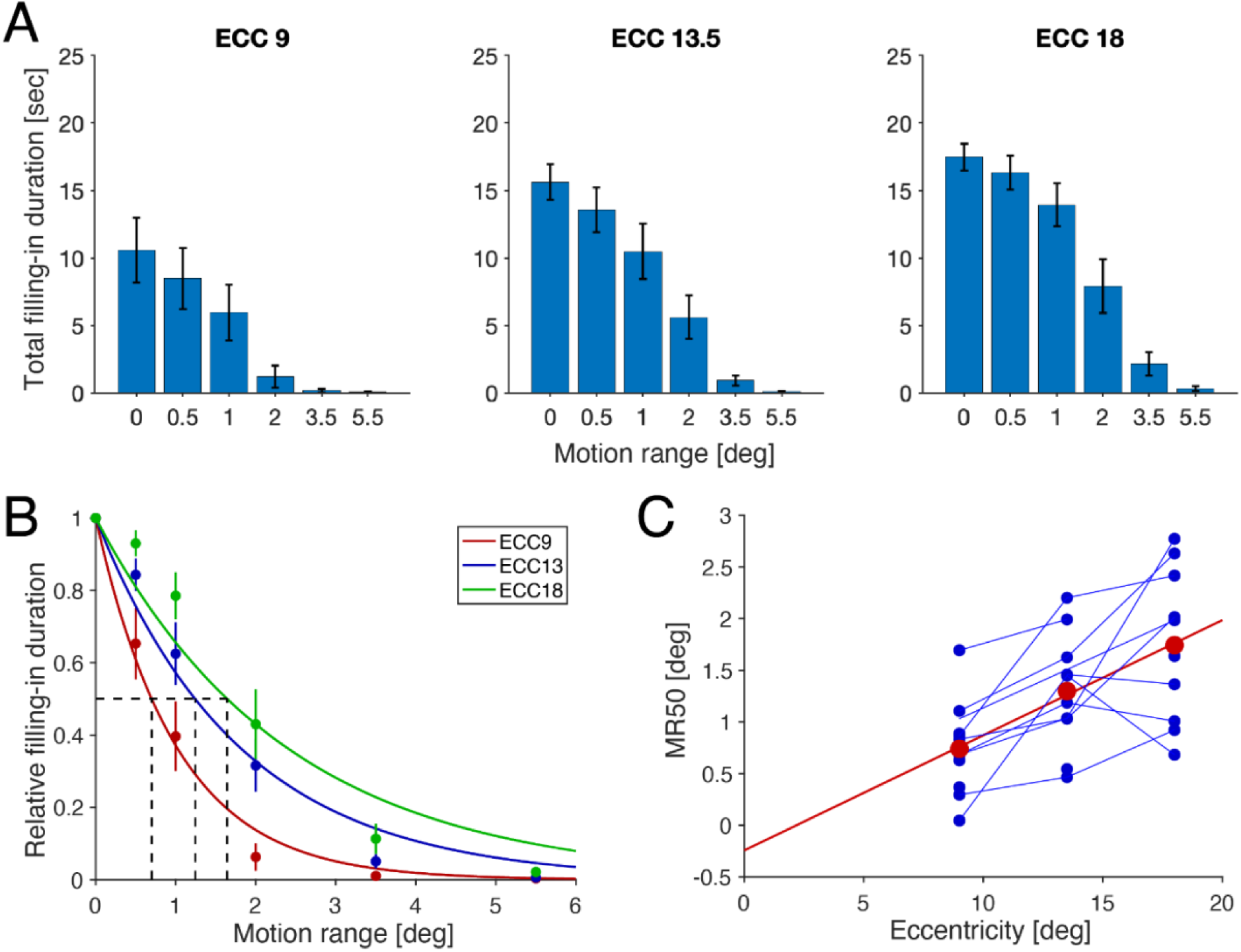
Mean total filling-in duration as a function of movement range, **A)** Averaged total filling-in duration over participants, error bars show standard error across participants. **B)** Mean total filling-in (Dots, vertical lines show standard error) at eccentricities 9°, 13.5° and 18°, normalized per eccentricity by the filling-in duration observed at the 0° motion range, and fitted with an exponential psychometric function. Dashed lines indicate the motion range associated with 50% of the maximum (MR50). **C)** MR50 per participant (blue dots) and of the sample mean (red dots) as a function of eccentricity. The red line represents a linear fit of the average MR50 plotted as a function of eccentricity. The blue dots are connected for participants who attended two or three sessions.

Mean normalized filling-in duration was plotted as a function of motion range per eccentricity in Fig. 2B (dots), and the data for each eccentricity were fitted with an exponential psychometric function (Fig. 2B lines, here fitted to the means over participants, see Methods for details). Using this function, fitted to the data per participant, we determined for each eccentricity and participant the MR50 (Fig. 2C blue dots), corresponding to the motion range permitting 50% of the filling-in duration observed in the no-motion condition. In Fig. 2C, we plotted individual participants’ MR50 as a function of eccentricity. A linear function fitted to the mean MR50 per eccentricity (Fig. 2C red dots) data yielded a slope of 0.11. This value compares well with the slope of the relationship between RF size and eccentricity in V1 and V2 (Klink et al., 2021, value of 0.0 - 0.21 in their figure 12; Wandell and Winawer, 2015, value of 0.15 estimated from their figure 1) but is outside of the range for higher areas (e.g. V4: Klink et al., 2021, value of 0.25 - 0.30 in their figure 12; Wandell and Winawer, 2015, value of 0.5 estimated from their figure 1).

We considered that the precision of our MR50 estimation may have been suboptimal at the 9° eccentricity since MR50 in this case (0.70°) fell close to the smallest motion range tested (0.5°). To verify our measurements at that eccentricity, we collected an additional session at 9° using a set of smaller motion ranges (0°, 0.33°, 0.67°, 1.33°, 2.33°, 3.67°). Data from this additional session with smaller motion ranges (Fig. 3B) were compared to the data collected with the standard set of motion ranges (Fig. 3A). When normalized total filling-in time (see Methods) was plotted as a function of motion size for the two sets of motion ranges (Fig. 3C), the fitted psychometric functions were very similar, and indeed there was no significant difference in the value of MR50 between the two sets (two-sample t-test, t(18) = 0.40, p = 0.69) (Fig. 3D). Here, we further calculated Bayes Factor (BF_01_ = 2.37). While acknowledging that the Bayes analysis only anecdotally supported the null hypothesis, given small sample in our data, this indicates that the MR50 estimate was robust against changes in the set of motion ranges used, supporting our approach for measuring motion dependence of perceptual filling-in.

**Figure 3.**
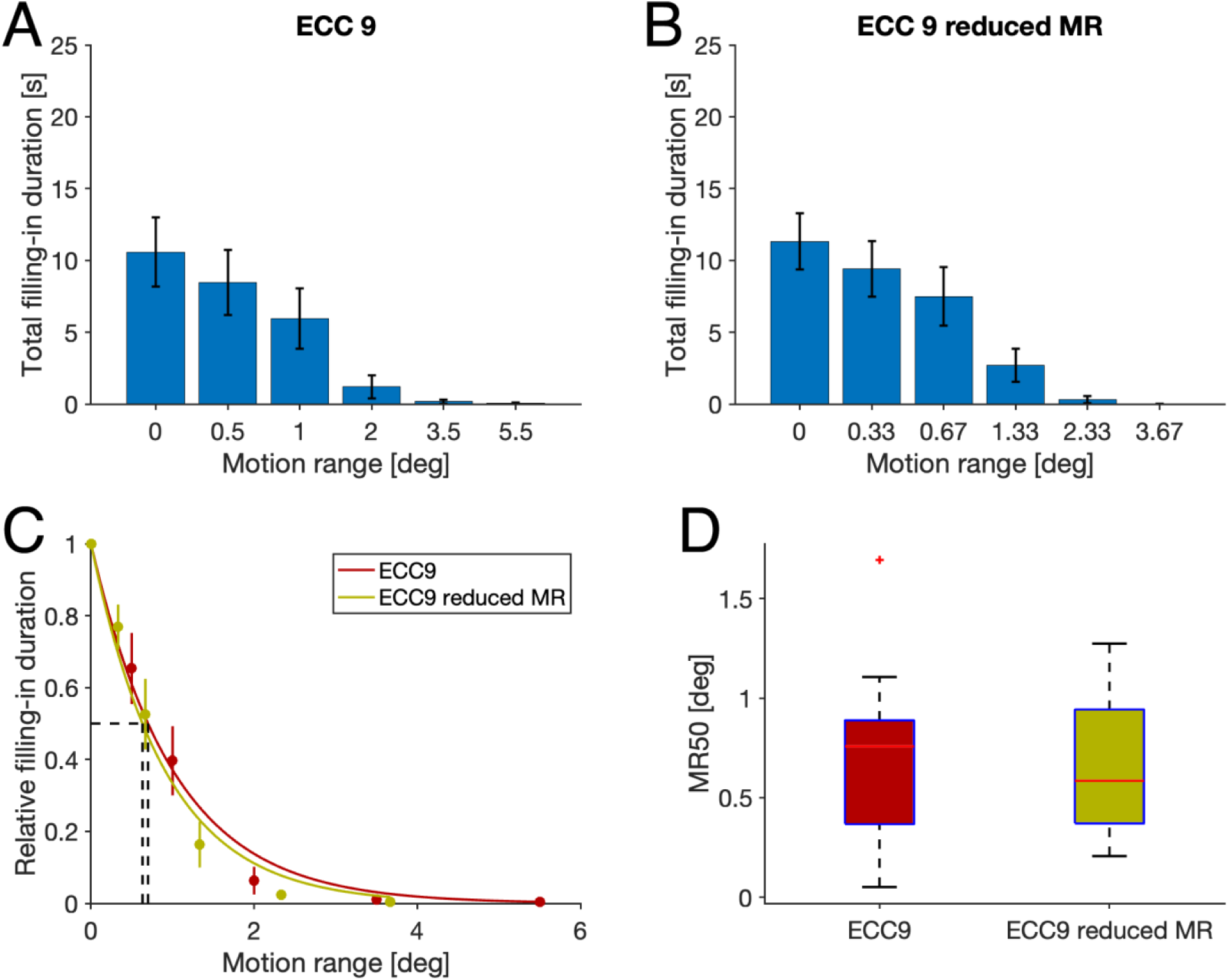
Control measurement of MR50 at 9° eccentricity using different ranges of motion. **A)** Total filling-in duration as a function of motion range in the restricted ranges of motion. **B)** As in A, for the standard motion range set, replotted from Figure 2A. **C)** Normalized filling-in duration as a function of motion range for both sets. Means and standard errors across participants are shown as dots and vertical bars. **D)** Boxplots of individual participants’ MR50 from the two datasets.

The results presented so far demonstrated a reduction of filling-in for larger motion ranges of the figure. While we interpret this in terms of the degree to which the figure boundaries stay within the RF of putative neurons, an alternative explanation may be that the larger motion ranges increased the rate of eye movements. Since MSs have been found to limit perceptual filling-in (Martinez-Conde et al., 2006; Troncoso et al., 2008), and since moving image elements have been found to induce eye movements (Gellman et al., 1990), this may be plausible. To test this possibility, we identified MSs from the eye movement data using the Engbert & Kliegl (2003) algorithm (see Method for more details). The majority (77%) of identified MSs were of less than 1° and 90% were below 1.5° (Fig. 4A). Further, the size and speed of MSs were highly correlated (Fig. 4A inset), in line with the ballistic nature of MSs (Martinez-Conde et al., 2004). To test whether MS characteristics differed across motion ranges and eccentricities, we compared the MS size and rate between the stimulus conditions (Fig. 4B). We found no main effect of motion range but an effect of eccentricity on MS size (a repeated measures ANOVA, interaction F(10, 150) = 0.13, p = 0.999; motion ranges F(5, 150) = 0.27, p = 0.932; eccentricities F(2, 150) = 0.53, p = 0.592). Bayesian ANOVA supported null hypothesis for interaction and motion range (interaction BF_01_ = 319; motion ranges BF_01_ = 33.1; eccentricities BF_01_ = 0.636). We found no main effect of motion range or eccentricity on MS rate (a repeated measures ANOVA, interaction F(10, 150) = 0.06, p = 0.999; motion ranges F(5, 150) = 0.11, p = 0.989; eccentricities F(2, 150) = 0.38, p = 0.685). Bayesian ANOVA supported null hypothesis for interaction and motion range (interaction BF_01_ = 551; motion ranges BF_01_ = 49.5; eccentricities BF_01_ = 0.641). Since Bayes Factor analysis did not support the null hypothesis for eccentricity, post-hoc Bayesian two-sample t-tests were conducted (MS size 9° vs. 13.5° BF_01_ = 0.246, 9° vs. 18° BF_01_ = 1.33, 13.5° vs. 18° BF_01_ = 1.80, MS rate 9° vs. 13.5° BF_01_ = 1.95, 9° vs. 18° BF_01_ = 4.49, 13.5° vs. 18° BF_01_ = 3.27). These results showed in favor of null hypothesis or no favor except favor in alternative hypothesis in MS size between 9° and 13.5°. We also confirmed the MS rate reduction before onset of filling-in (Martinez-Conde et al., 2006; Troncoso et al., 2008) using time-resolved cluster-based t-tests with 1000 permutations. Fig. 4C shows MS rate in a time window extending 5 sec before and after the start of the filling-in episode indicating significant MS rate drop from 0.75 sec before to 0.05 sec after filling-in onset. Taken together, these results confirm the relationship between MSs and filling-in but reject the possibility that this effect mediates the behavioral effects of motion range and mostly eccentricity described above.

**Figure 4.**
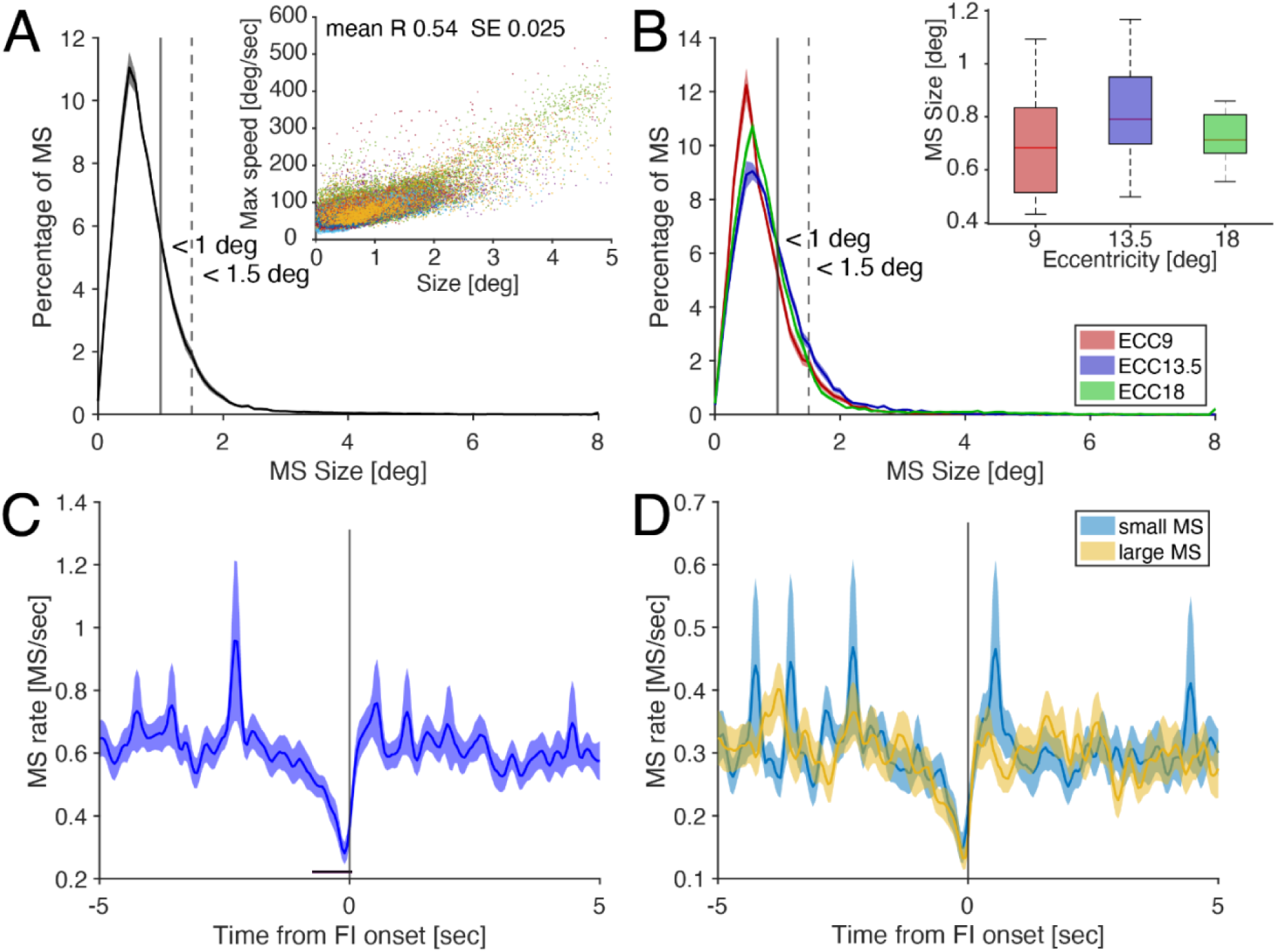
MSs and filling-in reports. **A)** Histogram of MS size averaged across participants. The shade shows standard error. Over all participants, 77% of MS were less than 1° and 90% of MS were less than 1.5°. **A inset)** relationship between MS size and MS max speed. Different color shows different participants. The correlation for all participants was high and positive value, mean of 0.54, SE of 0.025. **B)** Histogram of MS size averaged across participants at each eccentricity. Over all participants within same eccentricity sessions, 79% of MS were less than 1° and 91% of MS were less than 1.5° at 9° eccentricity, 69% of MS were less than 1° and 86% of MS were less than 1.5° at 13.5° eccentricity, and 76% of MS were less than 1° and 91% of MS were less than 1.5° at 18° eccentricity. **B inset)** Median of MS size of individual participants. **C)** MS rate time locked to filling-in onset in non-motion (0° motion) condition averaged across participants. Error bars show standard error. Black bar under the plot shows significant different cluster compared to other time windows. **D)** MS rate time locked to filling-in onset in non-motion condition after separating MS into two groups by its individual median MS size, averaged across participants. Error bars show standard error. Note that MS rate in this figure is half the rate as in panel B due to splitting MSs into two groups.

One interpretation of why MSs suppress perceptual filling-in is that the eye movement shifts the figure boundary on the retina, reducing adaptation, similar to our experimental manipulation of moving the figure on the screen. In this case, one could expect smaller MSs to be less effective in preventing filling-in than larger MSs, just as smaller movement ranges were less effective at reducing filling-in than larger motion ranges. Thus, one would expect the reduction in MSs prior to filling-in onset to be especially prominent for large MSs. To test this idea, we applied a median split to the distribution of MSs, to obtain ’small’ and ’large’ MSs for per session. Notably, the mean cutoff between small and large MSs was 0.72° with standard error of 0.029° across sessions, thus of a similar order magnitude of the MR50. We compared the frequency of these two classes of MSs in 0.05 sec time bins between 5 sec prior to and 5 sec following filling-in onset (Fig. 4D). For none of the time bins did we find a significant difference (paired t-tests, p > 0.05, corrected by cluster-based permutation test with 1000 permutations). We then calculated Bayes Factors in each time point. The median in all time point when we detect MS rate drop (0.75 sec before to 0.05 sec after filling-in onset) was BF_01_ = 5.01, which implies moderate evidence in favor of null hypothesis. Because the differential effect of small versus large MSs might differ across eccentricities, we repeated this analysis for data split over the three eccentricities. This analysis also showed no difference in the decrease of small and large MSs prior to filling-in at all three eccentricities (paired t-tests, p > 0.05, corrected by cluster-based permutation test with 1000 permutations for all three eccentricities).

## Discussion

In this study, we investigated perceptual filling-in of a moving figure in a Troxler paradigm. To our knowledge, this is the first study to parametrically investigate the effect of experimentally induced figure motion in this paradigm. Because boundary adaptation is considered to be a crucial component of perceptual filling-in, and because RF size could be considered an upper limit for figure movement that still allows boundary adaptation, we parametrically manipulated the motion range of the figure. We hypothesized that filling-in would be disrupted if the figure boundary moved beyond the size of a RF. By psychophysically measuring the motion range that hindered filling-in at several eccentricities, we obtained an estimate of the RF sizes involved in the boundary adaptation process at these different eccentricities. These estimates of RF size showed a linear relationship with eccentricity, with a slope consistent with how fMRI population RF estimates (Wandell and Winawer, 2015) and neurophysiological measurements of RFs (Klink et al., 2021) relate to eccentricity in V1 and V2, but not in higher areas. This supports the crucial involvement of low-level visual areas in the boundary adaptation that precedes filling-in in the Troxler paradigm. Importantly, MS rates did not differ among motion ranges or among eccentricities, which makes it plausible to investigate the efficiency of adaptation prior to filling-in in terms of the relation between retinal figure movement and neural RF sizes.

Some theoretical frameworks propose that the neurons that provide input to boundary representations (and that may be involved in the adaptation process) are also the neurons that provide input to a surface representation system in which activity related to the surface feature spreads laterally (Grossberg, 2003; Neumann et al., 2001; Paradiso et al., 1989). This suggests that the neural spreading of activity that underlies filling-in (Pessoa et al., 1998) may occur in the same areas where boundary adaptation takes place. In this view, our findings supporting V1 and V2 as a site for boundary adaptation, are consistent with several studies that have reported neural correlates of filling-in in early visual cortex. For example, the illusory brightness differences in a Craik-O’Brain-Cornsweet stimulus suggest a spreading of brightness initiated at the boundary, which according to neurophysiological data from (Hung et al., 2007; Roe et al., 2005) involves neural processes in V2 thin stripes. Neural correlates of surface completion also have been reported in V1 in fMRI studies using Kanizsa figures (Kok et al., 2016; Sasaki and Watanabe, 2004). Surfaces are also completed through the blind spot, and neurophysiological studies have found correlates of that in V1 (Fiorani Júnior et al., 1992; Komatsu et al., 2000). Furthermore, a neurophysiological correlate of perceptual filling-in a Troxler paradigm has been reported in monkey V2 and V3 (De Weerd et al., 1995). Nevertheless, the likely contributions of low-level areas to both neural adaptation effects underlying boundary adaptation and neural interpolation effects underlying perceptual filling-in do not exclude possible contributions of feedback (Roelfsema et al., 2002).

Previous studies of perceptual filling-in (Martinez-Conde et al., 2006; Troncoso et al., 2008), as well as our own data, show that filling-in episodes are preceded by a decline in MS rate. Here, we showed that figure movement amplitudes should remain within low-level neurons’ RFs to promote perceptual filling-in. If keeping figure-boundary movement within the confines of RF size is crucial to induce adaptation, the same factor may underlie the effectiveness of reduced MS rate in facilitating filling-in. In line with this reasoning, the decrease in the rate of larger MSs prior to filling-in might be predicted to be much larger than for smaller MSs, if larger MSs indeed were more efficient in preventing filling-in. Our data, however, did not support this prediction. Instead, the reduction in the rate of MS events seemed important in enabling filling-in, rather than the size of the MSs and the associated retinal displacements. Notably, the median of MS sizes in our data were somewhat smaller than the amplitudes of figure movements (MR50) that induced reduced filling-in. Therefore, the smaller MSs were not of sufficient size to expect their increased efficiency to prevent filling-in. However, these MSs still showed strong rate reduction before filling-in, suggesting that even small MSs can suppress perceptual filling-in. It may therefore be that MSs prevent boundary adaptation by mechanisms other than the bottom-up consequences of the retinal shifts of the stimulus. A plausible non-retinal mechanism that could underlie the prevention (or interruption) of filling-in could be a corollary discharge in visual cortex originating from the motor preparation of eye movements, fed back to the visual system to prepare it for impending retinal image shifts (Duhamel et al., 1992; Moore and Armstrong, 2003; Morris and Krekelberg, 2019; Sheliga et al., 1994). We speculate that such corollary discharge may reset both the adaptation process and any neural spreading processes underlying filling-in.

In our main experimental design, we used the same sized figure and the same set of motion ranges at the different eccentricities, with the same motion frequency (and therefore different motion speed) across different motion ranges. Since the properties of visual neurons change with eccentricity, our design therefore may have introduced unintended confounds. First, since we used the same figure size at all three eccentricities, the figure boundary representation was larger at the smaller eccentricity than at the greater eccentricity. Second, we adjusted the frequency of the motion to keep the number of potential boundary crossings constant across different motion ranges at a given eccentricity, thus, inevitably, there were motion speed differences across motion ranges. Given that motion induces saliency (Rosenholtz, 1999), the figure in our larger (and therefore faster) motion range conditions could have had higher saliency than ones in smaller motion range conditions. Third, motion speed for each motion range condition was identical across different eccentricities, which could introduce a confound as preferred neuronal motion speed tends to increase with eccentricity (Orban et al., 1986). However, neural speed tuning curves are typically quite broad (Beyeler et al., 2014; Maunsell and Van Essen, 1983; Orban et al., 1986; Priebe et al., 2006), making it unlikely that changes in eccentricity limited to 9-18° would have resulted in major differences in the sensitivity to the figure movement. Yet, small effects on our data cannot be excluded. Importantly, in a control experiment at one of the eccentricities, we found identical results using a reduced motion range at the smallest eccentricity, although the use of a reduced motion range entailed several of the subtle stimulus differences compared to the main experiment discussed above. This indicates that our methods were robust, and that the effects of possible stimulus confounds were small and did not affect in a major way the relationship we observed between estimated RF size and eccentricity.

In conclusion, our study is, to our knowledge, the first to parametrically investigate the effect of figure motion in a Troxler perceptual filling-in paradigm. Behavioral filling-in reports showed less filling-in for larger motion ranges of the figure, and more robust filling-in at larger eccentricities. Moreover, our analysis supported the hypothesis that boundary adaptation happens in early visual cortices like V1 and V2, and that the efficiency of adaptation relates to the magnitude of retinal shifts of the figure boundaries. An analysis of the effect of small and large MSs suggested that the retinal displacement *per se* caused by MSs may be insufficient to understand the facilitatory effects of MS rate decreases on filling-in. Instead, extra-retinal mechanisms may be at play as well. Our data confirm the relevance of low-level visual cortex for the adaptation process that enables perceptual filling-in. In addition, our data suggest that the effects of physical figure movements and MS on adaptation and filling-in are mediated through different neural mechanisms.

## References

Anstis, S., 2010. Visual filling-in. Curr. Biol. 20, R664–R666. 10.1016/j.cub.2010.06.029

Bart Krekelberg, 2024. Matlab Toolbox for Bayes Factor Analysis. 10.5281/ZENODO.13744717

Beyeler, M., Richert, M., Dutt, N.D., Krichmar, J.L., 2014. Efficient Spiking Neural Network Model of Pattern Motion Selectivity in Visual Cortex. Neuroinformatics 12, 435–454. 10.1007/s12021-014-9220-y

Bosman, C.A., Womelsdorf, T., Desimone, R., Fries, P., 2009. A Microsaccadic Rhythm Modulates Gamma-Band Synchronization and Behavior. J. Neurosci. 29, 9471–9480. 10.1523/JNEUROSCI.1193-09.2009

Corbett, J.J., Chen, J., 2018. The Visual System, in: Fundamental Neuroscience for Basic and Clinical Applications. Elsevier, pp. 286–305.e1. 10.1016/B978-0-323-39632-5.00020-7

De Weerd, P., 2006. Perceptual filling-in: more than the eye can see, in: Progress in Brain Research. Elsevier, pp. 227–245. 10.1016/S0079-6123(06)54012-9

De Weerd, P., Desimone, R., Ungerleider, L.G., 1998. Perceptual filling-in: a parametric study. Vision Res. 38, 2721–2734. 10.1016/S0042-6989(97)00432-X

De Weerd, P., Gattass, R., Desimone, R., Ungerleider, L.G., 1995. Responses of cells in monkey visual cortex during perceptual filling-in of an artificial scotoma. Nature 377, 731–734. 10.1038/377731a0

Duhamel, J.-R., Colby, C.L., Goldberg, M.E., 1992. The Updating of the Representation of Visual Space in Parietal Cortex by Intended Eye Movements. Science 255, 90–92. 10.1126/science.1553535

Dumoulin, S.O., Wandell, B.A., 2008. Population receptive field estimates in human visual cortex. NeuroImage 39, 647–660. 10.1016/j.neuroimage.2007.09.034

Engbert, R., Kliegl, R., 2003. Microsaccades uncover the orientation of covert attention. Vision Res. 43, 1035–1045. 10.1016/S0042-6989(03)00084-1

Fiorani Júnior, M., Rosa, M.G., Gattass, R., Rocha-Miranda, C.E., 1992. Dynamic surrounds of receptive fields in primate striate cortex: a physiological basis for perceptual completion? Proc. Natl. Acad. Sci. 89, 8547–8551. 10.1073/pnas.89.18.8547

Gattass, R., Gross, C.G., Sandell, J.H., 1981. Visual topography of V2 in the macaque. J. Comp. Neurol. 201, 519–539. 10.1002/cne.902010405

Gattass, R., Sousa, A., Gross, C., 1988. Visuotopic organization and extent of V3 and V4 of the macaque. J. Neurosci. 8, 1831–1845. 10.1523/JNEUROSCI.08-06-01831.1988

Gattass, R., Sousa, A.P.B., Rosa, M.G.P., 1987. Visual topography of V1 in the *Cebus* monkey. J. Comp. Neurol. 259, 529–548. 10.1002/cne.902590404

Gellman, R.S., Carl, J.R., Miles, F.A., 1990. Short latency ocular-following responses in man. Vis. Neurosci. 5, 107–122. 10.1017/S0952523800000158

Greilich, J., Baumann, M.P., Hafed, Z.M., 2024. Microsaccadic suppression of peripheral perceptual detection performance as a function of foveated visual image appearance. J. Vis. 24, 3. 10.1167/jov.24.11.3

Grossberg, S., 2003. Filling-in the forms: Surface and boundary interactions in visual cortex. Oxford University Press. 10.1093/acprof:oso/9780195140132.001.0001

Grossberg, S., Todorovic, D., 1988. Neural dynamics of 1-D and 2-D brightness perception: A unified model of classical and recent phenomena. Percept. Psychophys. 43, 241–277. 10.3758/BF03207869

Herrington, T.M., Masse, N.Y., Hachmeh, K.J., Smith, J.E.T., Assad, J.A., Cook, E.P., 2009. The Effect of Microsaccades on the Correlation between Neural Activity and Behavior in Middle Temporal, Ventral Intraparietal, and Lateral Intraparietal Areas. J. Neurosci. 29, 5793–5805. 10.1523/JNEUROSCI.4412-08.2009

Hubel, D.H., Wiesel, T.N., 1959. Receptive fields of single neurones in the cat’s striate cortex. J. Physiol. 148, 574–591. 10.1113/jphysiol.1959.sp006308

Hung, C.P., Ramsden, B.M., Roe, A.W., 2007. A functional circuitry for edge-induced brightness perception. Nat. Neurosci. 10, 1185–1190. 10.1038/nn1948

Hunzelmann, N., Spillmann, L., 1984. Movement adaptation in the peripheral retina. Vision Res. 24, 1765–1769. 10.1016/0042-6989(84)90007-5

Jeffreys, H., 1998. The theory of probability, 3rd edition. ed. Oxford University Press.

Klink, P.C., Chen, X., Vanduffel, W., Roelfsema, P.R., 2021. Population receptive fields in nonhuman primates from whole-brain fMRI and large-scale neurophysiology in visual cortex. eLife 10. 10.7554/elife.67304

Kok, P., Bains, L.J., van Mourik, T., Norris, D.G., de Lange, F.P., 2016. Selective Activation of the Deep Layers of the Human Primary Visual Cortex by Top-Down Feedback. Curr. Biol. 26, 371–376. 10.1016/j.cub.2015.12.038

Komatsu, H., Kinoshita, M., Murakami, I., 2000. Neural Responses in the Retinotopic Representation of the Blind Spot in the Macaque V1 to Stimuli for Perceptual Filling-In. J. Neurosci. 20, 9310–9319. 10.1523/jneurosci.20-24-09310.2000

Lima, B., Florentino, M.M., Fiorani, M., Soares, J.G.M., Schmidt, K.E., Neuenschwander, S., Baron, J., Gattass, R., 2023. Cortical maps as a fundamental neural substrate for visual representation. Prog. Neurobiol. 224, 102424. 10.1016/j.pneurobio.2023.102424

Lowet, E., Roberts, M.J., Bosman, C.A., Fries, P., De Weerd, P., 2016. Areas V1 and V2 show microsaccade-related 3–4-Hz covariation in gamma power and frequency. Eur. J. Neurosci. 43, 1286–1296. 10.1111/ejn.13126

Martinez-Conde, S., Macknik, S.L., Hubel, D.H., 2004. The role of fixational eye movements in visual perception. Nat. Rev. Neurosci. 5, 229–240. 10.1038/nrn1348

Martinez-Conde, S., Macknik, S.L., Troncoso, X.G., Dyar, T.A., 2006. Microsaccades Counteract Visual Fading during Fixation. Neuron 49, 297–305. 10.1016/j.neuron.2005.11.033

Matin, E., 1974. Saccadic suppression: A review and an analysis. Psychol. Bull. 81, 899–917. 10.1037/h0037368

Maunsell, J.H., Van Essen, D.C., 1983. Functional properties of neurons in middle temporal visual area of the macaque monkey. I. Selectivity for stimulus direction, speed, and orientation. J. Neurophysiol. 49, 1127–1147. 10.1152/jn.1983.49.5.1127

Moore, T., Armstrong, K.M., 2003. Selective gating of visual signals by microstimulation of frontal cortex. Nature 421, 370–373. 10.1038/nature01341

Morris, A.P., Krekelberg, B., 2019. A Stable Visual World in Primate Primary Visual Cortex. Curr. Biol. 29, 1471–1480.e6. 10.1016/j.cub.2019.03.069

Neumann, H., Pessoa, L., Hansen, T., 2001. Visual filling-in for computing perceptual surface properties. Biol. Cybern. 85, 355–369. 10.1007/s004220100258

Nyström, M., Andersson, R., Holmqvist, K., Van De Weijer, J., 2013. The influence of calibration method and eye physiology on eyetracking data quality. Behav. Res. Methods 45, 272–288. 10.3758/s13428-012-0247-4

Orban, G.A., Kennedy, H., Bullier, J., 1986. Velocity sensitivity and direction selectivity of neurons in areas V1 and V2 of the monkey: influence of eccentricity. J. Neurophysiol. 56, 462–480. 10.1152/jn.1986.56.2.462

Paradiso, M.A., Nakayama, K., 1991. Brightness perception and filling-in. Vision Res. 31, 1221–1236. 10.1016/0042-6989(91)90047-9

Paradiso, M.A., Shimojo, S., Nakayama, K., 1989. Subjective contours, tilt aftereffects, and visual cortical organization. Vision Res. 29, 1205–1213. 10.1016/0042-6989(89)90066-7

Pessoa, L., Thompson, E., Noë, A., 1998. Finding out about filling-in: A guide to perceptual completion for visual science and the philosophy of perception. Behav. Brain Sci. 21, 723–748. 10.1017/S0140525X98001757

Priebe, N.J., Lisberger, S.G., Movshon, J.A., 2006. Tuning for Spatiotemporal Frequency and Speed in Directionally Selective Neurons of Macaque Striate Cortex. J. Neurosci. 26, 2941–2950. 10.1523/JNEUROSCI.3936-05.2006

Ramachandran, V.S., Gregory, R.L., Aiken, W., 1993. Perceptual fading of visual texture borders. Vision Res. 33, 717–721. 10.1016/0042-6989(93)90191-X

Roe, A.W., Lu, H.D., Hung, C.P., 2005. Cortical processing of a brightness illusion. Proc. Natl. Acad. Sci. 102, 3869–3874. 10.1073/pnas.0500097102

Roelfsema, P.R., Lamme, V.A.F., Spekreijse, H., Bosch, H., 2002. Figure—Ground Segregation in a Recurrent Network Architecture. J. Cogn. Neurosci. 14, 525–537. 10.1162/08989290260045756

Rosenholtz, R., 1999. A simple saliency model predicts a number of motion popout phenomena. Vision Res. 39, 3157–3163. 10.1016/s0042-6989(99)00077-2

Sasaki, Y., Watanabe, T., 2004. The primary visual cortex fills in color. Proc. Natl. Acad. Sci. 101, 18251–18256. 10.1073/pnas.0406293102

Sheliga, B.M., Riggio, L., Rizzolatti, G., 1994. Orienting of attention and eye movements. Exp. Brain Res. 98. 10.1007/BF00233988

Sincich, L.C., Horton, J.C., 2005. THE CIRCUITRY OF V1 AND V2: Integration of Color, Form, and Motion. Annu. Rev. Neurosci. 28, 303–326. 10.1146/annurev.neuro.28.061604.135731

Sperry, R.W., 1950. Neural basis of the spontaneous optokinetic response produced by visual inversion. J. Comp. Physiol. Psychol. 43, 482–489. 10.1037/h0055479

Spillmann, L., Kurtenbach, A., 1992. Dynamic noise backgrounds facilitate target fading. Vision Res. 32, 1941–1946. 10.1016/0042-6989(92)90053-L

Troncoso, X.G., Macknik, S.L., Martinez-Conde, S., 2008. Microsaccades counteract perceptual filling-in. J. Vis. 8, 15–15. 10.1167/8.14.15

Troxler, D., 1804. Über das Verschwinden gegebener Gegenstände innerhalb unseres Gesichtskreises. Ophthalmol. Bibl.

Wandell, B.A., Winawer, J., 2015. Computational neuroimaging and population receptive fields. Trends Cogn. Sci. 19, 349–357. 10.1016/j.tics.2015.03.009

Weil, R.S., Rees, G., 2011. A new taxonomy for perceptual filling-in. Brain Res. Rev. 67, 40–55. 10.1016/j.brainresrev.2010.10.004

Welchman, A.E., Harris, J.M., 2001. Filling-in the details on perceptual fading. Vision Res. 41, 2107–2117. 10.1016/S0042-6989(01)00087-6

Wiesel, T.N., Hubel, D.H., 1963. EFFECTS OF VISUAL DEPRIVATION ON MORPHOLOGY AND PHYSIOLOGY OF CELLS IN THE CAT’S LATERAL GENICULATE BODY. J. Neurophysiol. 26, 978–993. 10.1152/jn.1963.26.6.978

Zuber, B.L., Stark, L., 1966. Saccadic suppression: Elevation of visual threshold associated with saccadic eye movements. Exp. Neurol. 16, 65–79. 10.1016/0014-4886(66)90087-2

